# VolPy: automated and scalable analysis pipelines for voltage imaging datasets

**DOI:** 10.1101/2020.01.02.892323

**Authors:** Changjia Cai, Johannes Friedrich, Eftychios A Pnevmatikakis, Kaspar Podgorski, Andrea Giovannucci

**Affiliations:** Joint Department of Biomedical Engineering at University of North Carolina at Chapel Hill and North Carolina State University, Chapel Hill, NC, USA; Flatiron Institute, Simons Foundation, New York, NY, USA; Janelia Research Campus, Howard Hughes Medical Institute, Ashburn, VA, USA; Neuroscience Center, University of North Carolina at Chapel Hill, Chapel Hill, NC, USA

## Abstract

Voltage imaging enables monitoring neural activity at sub-millisecond and sub-compartment scale, and therefore opens the path to studying sub-threshold activity, synchrony, and network dynamics with unprecedented spatio-temporal resolution. However, high data rates (>800MB/s) and low signal-to-noise ratios have created a severe bottleneck for analysis of such datasets. Here we present *VolPy*, the first turn-key, automated and scalable pipeline to pre-process voltage imaging datasets. *VolPy* features fast motion correction, memory mapping, segmentation, and spike inference, all built on a highly parallelized and computationally efficient framework that optimizes memory and speed. Given the lack of single cell voltage imaging ground truth examples, we introduce a corpus of 24 manually annotated datasets from different preparations and voltage indicators. We benchmark *VolPy* against this corpus and electrophysiology recordings, demonstrating excellent performance in neuron localization, spike extraction, and scalability.

## Introduction

While several methods have been developed to process voltage imaging data at mesoscopic scale and multi-unit resolution (***Marshall et al., 2016***; ***Carandini et al., 2015***; ***Akemann et al., 2012***), to date there is no established pipeline for large-scale single cell analysis, which was only recently necessitated by sensitive new voltage indicators (***Knöpfel and Song, 2019***; ***Abdelfattah et al., 2019***; ***Adam et al., 2019***; ***Kannan et al., 2018***; ***Piatkevich et al., 2019***, ***2018***; ***Roome and Kuhn, 2018***). Indeed, voltage imaging datasets present significant new challenges compared to calcium imaging, calling for new approaches. On the one hand, dataset sizes have increased one or two orders of magnitude (Tens of GBs vs TBs per hour), and on the other hand, assumptions of existing calcium imaging analysis methods may be inappropriate. For instance, non-negative matrix factorization (NMF) methods (***Giovannucci et al., 2019***) fail when applied to voltage imaging data for three reasons (***Buchanan et al., 2018***): (i) while good segmentation approaches exist for somatic imaging, these fail for other imaging modalities, (ii) it is difficult to separate weak components from noise using current NMF approaches; (iii) since voltage traces typically display both positive and negative fluctuations around the baseline resting potential, the NMF framework, based on non-negativity in both spatial and temporal domains, is not readily applicable to voltage imaging data.

### Related work

Some relevant methods are beginning to populate the literature. For instance, ad-hoc solutions presented in (***Abdelfattah et al., 2019***) provide interesting starting points to extract and denoise spikes semi-automatically, but suffer from some drawbacks. First, they require manual or semi-manual selection of neurons, which is both labor intensive and prone to irreproducibility. Second, the algorithms do not scale well in computational time and memory. Finally, these algorithms are not embedded into a reusable and well documented format, which hinders their reuse by a broad community (***Teeters et al., 2015***). A more standardiffied approach is provided by (***Adam et al., 2019***; ***Buchanan et al., 2018***). However, this method does not embed an adaptive and automated mechanism for spike extraction and is not integrated in a robust, scalable and multi-platform framework. Further, lack of ground truth datasets has so far hindered the validation of all these approaches. In summary, no validated, complete, scalable and automatic analysis pipeline for voltage imaging data analysis exists to date.

### Contributions

To address these shortcomings, we established objective performance evaluation benchmarks and a new analysis pipeline for pre-processing voltage imaging data, which we named *VolPy*. First, in order to establish a common validation framework and to automate neuron segmentation, we created a corpus of annotated datasets with manually segmented neurons. Second, we used this benchmark to train a supervised algorithm to automatically localize and segment cells via convolutional networks (***He et al., 2017***). Third, we introduced an improved algorithm to denoise fluorescence traces and extract single spikes, which builds upon the SpikePursuit prototype (***Abdelfattah et al., 2019***). We modified the core SpikePursuit algorithm to achieve better performance and scalability, both by speeding up the underlying optimization algorithm, and by building the infrastructure to parallelize it efficiently and with low memory requirements. Notably, the algorithm is automatically initialized using the neural network for localizing and segmenting neurons, a task that was previously performed manually. Fourth, we quantitatively evaluated *VolPy* neuron segmentation, spike extraction and scalability. Segmentation was evaluated on 24 datasets, encompassing different brain areas, animal preparations and voltage indicators (Tables 1 and 2). The performance of *VolPy* on the validation set was high for datasets with more training samples, but progressively degraded when less data was available. When compared with electrophysiology data, *VolPy* spike extraction featured *F*_1_ scores mostly above 90% on three example neurons. The computational performance of *VolPy* was evaluated on the largest dataset available to us and showed promising results in terms of computational time (up to 66 frames/sec) and memory requirements (down to 1.5X RAM of the original dataset size).

**Table 1.** Properties of three heterogeneous types of datasets. For each type of dataset the name, organism, brain region, source, imaging rate, voltage indicator, and the total number of neurons selected by the manual annotators are given.

| Name | Organism | Brain region | Source | Rate (Hz) | Indicator | # neurons |
| --- | --- | --- | --- | --- | --- | --- |
| L1 | Mouse | L1 cortex | *Abdelfattah et al. (2019)* | 400 | Voltron | 523 |
| TEG | Zebrafish | Tegmental | *Abdelfattah et al. (2019)* | 300 | Voltron | 107 |
| HPC | Mouse | Hippocampus | *Adam et al. (2019)* | 1000 | paQuasAr3-s | 41 |

**Table 2.** All annotated datasets for segmentation of *VolPy*. For each dataset the name, size of datasets and number of neurons.

| Name | Size | # | Name | Size | # |
| --- | --- | --- | --- | --- | --- |
| L1.00.00 | 20000*512*128 | 84 | HPC.29.04 | 20000*164*96 | 2 |
| L1.01.00 | 20000*512*128 | 53 | HPC.29.06 | 20000*228*96 | 2 |
| L1.01.35 | 20000*512*128 | 69 | HPC.32.01 | 20000*256*96 | 4 |
| L1.02.00 | 20000*512*128 | 61 | HPC.38.05 | 20000*176*92 | 4 |
| L1.02.80 | 20000*512*128 | 43 | HPC.38.03 | 20000*128*88 | 2 |
| L1.03.00 | 20000*512*128 | 79 | HPC.39.07 | 20000*264*96 | 5 |
| L1.03.35 | 20000*512*128 | 57 | HPC.39.03 | 20000*276*96 | 5 |
| L1.04.00 | 20000*512*128 | 43 | HPC.39.04 | 20000*336*96 | 4 |
| L1.04.50 | 20000*512*128 | 34 | HPC.48.01 | 20000*224*96 | 2 |
| TEG.01.02 | 10000*364*320 | 33 | HPC.48.05 | 20000*212*96 | 4 |
| TEG.02.01 | 10000*360*256 | 29 | HPC.48.07 | 20000*280*96 | 2 |
| TEG.03.01 | 10000*508*288 | 45 | HPC.48.08 | 20000*284*96 | 3 |

We integrated our methods within the *CaImAn* ecosystem (***Giovannucci et al., 2019***), a popular suite of tools for single cell resolution brain imaging analysis. This integration allowed us to use and extend *CaImAn*’s tools for motion correction and memory mapping to enable scalability of our algorithms. In particular, we adapted *CaImAn* to perform motion correction (***Pnevmatikakis and Giovannucci, 2017***), memory mapping (***Giovannucci et al., 2019***), and run the modified SpikePursuit algorithm on voltage imaging data. Besides the obvious computational advantages, this made *VolPy* immediately available to the research labs already relying on the *CaImAn* ecosystem.

In summary, we have developed a validated, scalable, turn-key, documented and easily installed voltage imaging analysis pipeline that has been packaged into a popular open source software suite. This will enable an increasing number of laboratories to exploit the advantages provided by voltage imaging and therefore accelerate the pace of discovery in neuroscience.

The paper is organized as follows. We first report the new methods developed in *VolPy*, then we benchmark their performance, and finally we discuss some implications. We leave most of the fine implementation details for the Material and Methods section.

## Methods

### Creation of a corpus of annotated datasets

To date there is no metric to establish whether voltage imaging algorithms for single cell localization and/or segmentation perform well in practice. To overcome this problem, and with the goal of developing new supervised algorithms, we generated a corpus of annotated datasets (Ground truth, GT) in which neurons are manually segmented. GT is constructed by human labelers from two summary images (mean and local correlation images, Figure 1 B and C) and a pre-processed movie that highlights active neurons (local correlation movie, Suppl Movie 1). More specifically, after motion correction, we generate a mean image, a correlation image and a correlation video as follows:

*Mean image*. To compute the mean image, we average the movie across time for each pixel and normalize by the pixel-wise z-score.
*Correlation image*. The correlation image is a variation of that implemented in (***Smith and Häusser, 2010***), which is applied to a baseline-subtracted movie. To estimate the baseline of the movie, frames are first binned according to the window length (a parameter set to 1 second). We compute the 8^*th*^ running percentile of the signal for each pixel. Intermediate values of the baseline are inferred by spline interpolation, which is a fast approximation of a running window. After removing the baseline of the movie, we compute the correlation image of the movie by averaging the temporal correlation of each pixel with its eight neighbor pixels. We also normalize the correlation image by z-scoring when fed to the neural network.
*Correlation movie*. We introduce a novel type of denoising operation, the correlation movie. The correlation movie is essentially a running version of the correlation image computed over overlapping chunks of video frames. This new type of denoising significantly improves the visibility of spikes in voltage imaging movies (see Movie 1). There are two parameters governing the creation of the correlation movie, the chunk size (number of frames over which each correlation image is computed) and stride (the number of frames to skip between consecutive chunks).

**Figure 1.**
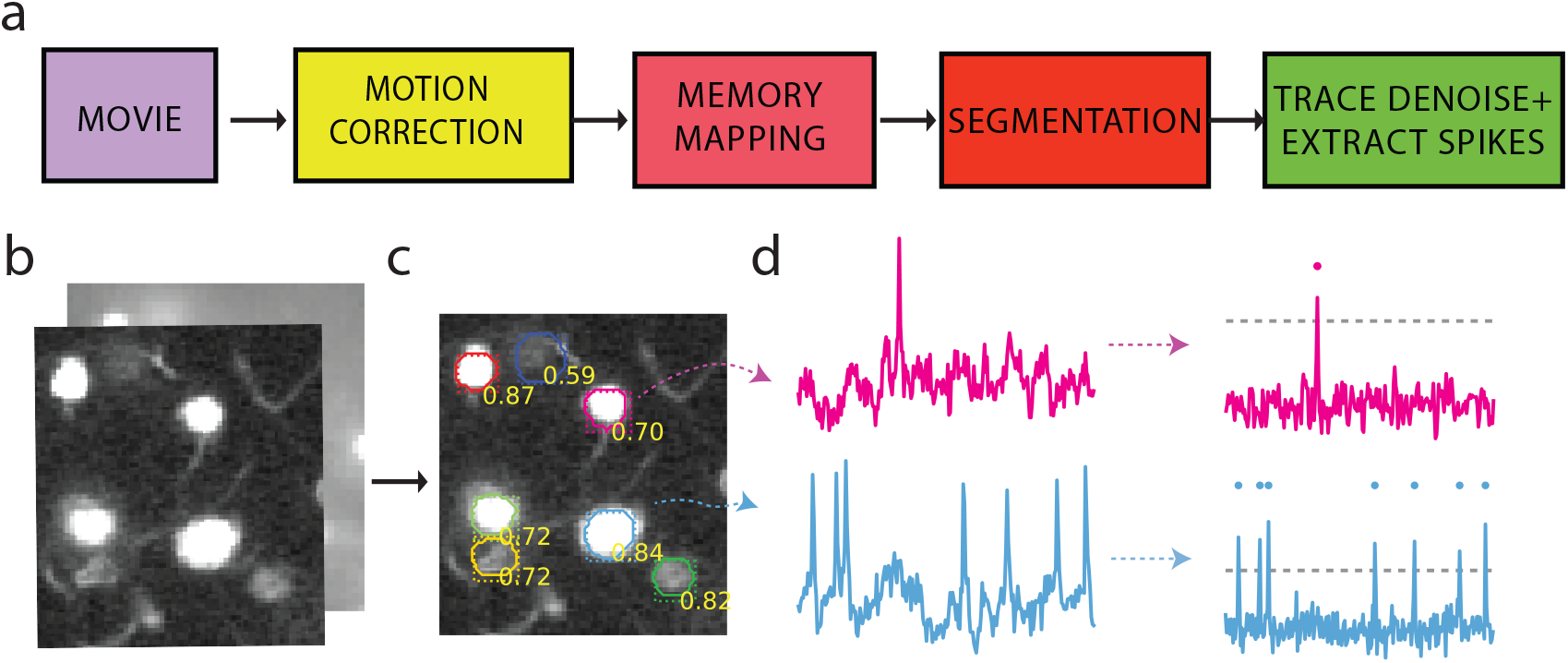
Analysis pip eline for voltage imaging data. (a) Four pre-processing steps are required to extract spikes and neuron locations from voltage imaging movies. (b) Correlation image (front) and mean image (back) of one of the Layer1 neocortex movies as the input of the segmentation step. (c) The segmentation step outputs class probabilities, bounding boxes and contours. The results are overlaid to the correlation image in (b). (d) Result of trace denoising and spike extraction. The gray dashed horizontal line represents the inferred spike threshold.

We implemented parallelized routines which allow to compute efficiently summary images and correlation movies. These routines need only to load in memory small contiguous chunks of the input movies and can process them efficiently in parallel over multiple cores.

Guided by these three visual cues, two annotators marked the contours of neurons using the ImageJ Cell Magic Wand tool plugin (***Walker, 2014***). For neurons to be selected both annotators had to agree on the selection, which had to fulfill the following criteria: (i) neurons were very clear on at least one of the three cues; (ii) Neurons were moderately clear in one of the summary images and exhibited a spatial footprint in selected frames of the local correlations movie (see Figure 8). Summary information about the annotated datasets is reported in Table 1. Examples of manual annotations are reported in Figure 3 (red contours).

### A novel analysis pipeline for voltage imaging

Voltage imaging is characterized by high data rates (up to 800 MB/sec). This often leads to the creation of movies that are difficult to manage using conventional computers. Even though scalable algorithms for calcium imaging exist (***Giovannucci et al., 2019***), they fail when applied to voltage imaging. Here we propose a novel scalable pipeline for automated analysis that performs preprocessing steps required to extract spikes and sub-threshold activity from voltage imaging movies. In Figure 1 we illustrate the proposed standard pipeline for analyzing voltage imaging data. First, input data is processed to remove motion artifacts with parallelized algorithms, and saved into a memory map file format that enables efficient concurrent access. In a second stage, *VolPy* localizes candidate neurons using supervised algorithms (Figure 1a and c). Finally, *VolPy* denoises fluorescence traces, infers spatial footprints, and extracts neural activity of each neuron through unsupervised learning (Figure 1a and d). Notice that the presented framework is modular, and therefore allows for easy testing of new algorithms by replacing individual components of the pipeline. In what follows we present each stage of the *VolPy* pipeline in detail.

#### Motion correction and memory mapping

First, movies need to be corrected for sample movement. We performed this registration relying on a variation of the algorithm described in (***Giovannucci et al., 2019***; ***Pnevmatikakis and Giovannucci, 2017***), which exploits multi-core parallelization and memory mapping to register frames to a template based on cross-correlation. The only variation with respect the original algorithm is that the new implementation can perform motion correction on a large number of small files containing a single image (a typical output format of fast imaging cameras). This avoids the memory-intensive job of transforming single image files into multi-page files, and limitations of file size. Motion correction, similarly to (***Giovannucci et al., 2019***), is performed in parallel over multiple segments of the same movie and the result is directly stored in a memory mapped file that is efficiently readable frame-by frame (Fortran (F) order, see Materials and Methods). Relying on the algorithms of *CaImAn*, we then efficiently create a second copy of the file that allows rapid pixel by pixel reads (C order, see Materials and Methods) instead of frame by frame (memory mapping, Figure 1a). This enables a fundamental feature of *VolPy*, that is the ability to quickly read arbitrary portions of the field of view in any direction without having to load the full movie into memory. In summary, the first two steps of the pipeline generate two copies of the motion corrected movie, one efficiently and concurrently read frame-by-frame, and one pixel by pixel. This allows parallelization of all the operations which are required to generate summary images and denoise the signal, as specified below.

#### Segmentation

The low SNR of voltage imaging data hinders the applicability of the segmentation methods previously devised for calcium imaging data (***Pnevmatikakis et al., 2016***). Here we propose to initialize denoising algorithms with supervised learning approaches. While previous attempts at cell localization and segmentation have extended U-Net fully convolutional network architectures (***Falk et al., 2019***), in our hands this family of methods failed when facing datasets in which neurons overlap (Figure 3a). We hypothesize that this happens since U-Net is a semantic segmentation approach, which aims at separating neurons pixels from the background pixels, and therefore performs poorly in our instance segmentation task of separating overlapping neurons. We approached the problem with Mask R-CNN, a convolutional network for object localization and segmentation (***He et al., 2017***). Mask R-CNN is a particularly promising architecture as it enables to separate overlapping objects in a specific area by providing each object with a unique bounding box.

The network, which is trained with a corpus of annotated datasets generated by us, takes summary images as input and outputs contours and bounding boxes of candidate neurons (Figure 1c), along with a class probability. An example of the network inference on a validation dataset by *VolPy* is shown in Figure 2. The resulting network performs well in our task on widely different datasets.

**Figure 2.**
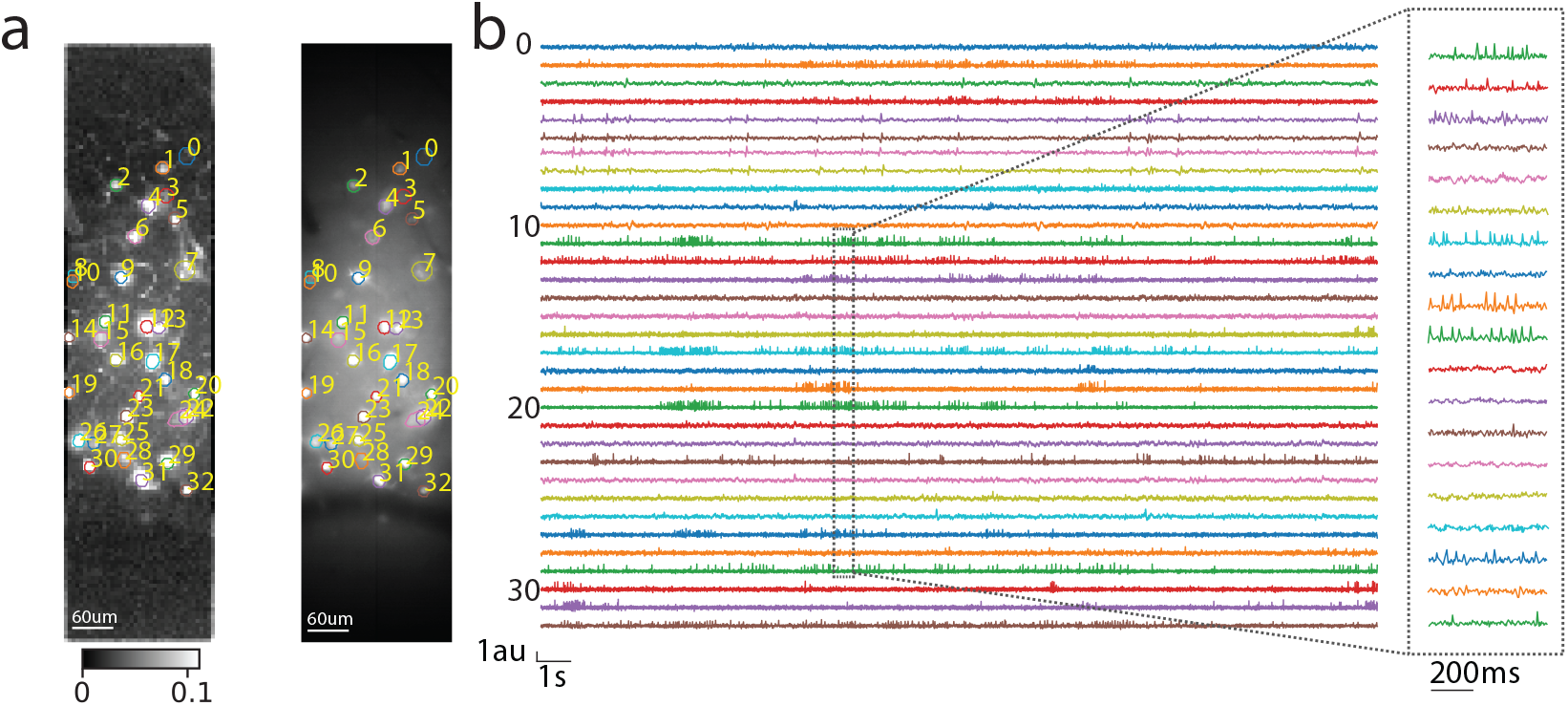
Result of processing a mouse L1 neocortex voltage imaging dataset using the *VolPy* pipeline. (a) Correlation image (left) and mean image (right) overlaid with contours detected by *VolPy*. (b) Temporal traces corresponding to neurons in panel (a) extracted by *VolPy* (left). The dashed gray portion of the traces is magnified on the right.

#### Trace denoising and spike extraction

Classical algorithms for denoising calcium imaging movies and extracting spikes from the corresponding fluorescence traces fail when applied to voltage imaging movies. On the one hand, the low signal-to-noise ratio and the complex background fluorescence require new methods for refining spatial footprints, and on the other hand, substantially different biophysical models underlie the temporal dynamics of the fluorescence associated to spikes. To solve both problems, we build upon and extend the SpikePursuit algorithm (***Abdelfattah et al., 2019***). In particular, we improve SpikePursuit in the following directions (see Material and Methods for details):

- While the original version of the algorithm required manual selection of candidate neurons, *VolPy* automatically initializes it using the output of the trained Mask R-CNN (Figure 1c and d).
- A minimal amount of data needs to be loaded in memory thanks to the memory mapping infrastructure, thereby reducing memory requirements.
- We increase the reliability of the underlying inference algorithm, by introducing a more robust estimate of the background.
- We scale up the performance by improving the algorithms which perform Ridge regression during inference of spikes and spatial masks.

#### Embarrassingly parallel computing in *VolPy*

Unlike *CaImAn*, which is based on a Map-Reduce framework to parallelize execution, *VolPy* relies on an embarrassingly parallel paradigm (***Herlihy and Shavit, 2011***). Embarrassingly parallel solutions exploit the lack of dependence among tasks to efficiently deploy concurrency. Indeed the core of *VolPy* algorithms decouples computations so that each neuron is processed independently.

First, motion correction in *VolPy* is parallelized by processing temporal chunks of movie data on different CPUs while saved in a memory mapped file which is efficiently read frame-by-frame. second, the various summary images and correlation movies can be computed in parallel processing contiguous temporal chunks of the memory mapped movies. Subsequently, the motion corrected file is processed and saved into another memory mapped file which efficiently read pixel-by-pixel. Finally, during trace denoising and spike extraction, candidate neurons can be processed in parallel without significant memory overhead based on the fact that the signal of each neuron is localized in pixels near to the center of the neuron. Exploiting this locality, *VolPy* processes in parallel context regions surrounding each candidate neuron (see Materials and Methods) by reading concurrently from the pixel-by-pixel memory mapped file. Each process extracts denoised fluorescence signals and spikes from the corresponding context region. In conclusion, *VolPy* enables automatic analysis of large scale voltage imaging datasets. In Figure 2, we report the result of preprocessing an example mouse L1 neocortex voltage imaging dataset with the *VolPy* pipeline.

## Results

In what follows we report a systematic evaluation of *VolPy* against ground truth in terms of performance in identifying neurons, spike extraction and scalability.

### *VolPy* localizes neurons using a moderate amount of training data

We trained a modified version of the Mask R-CNN network architecture (see Material and Methods for details) on three heterogeneous types of datasets (Table 1) and evaluated its performance using 3-fold cross validation (see Table 2 and Materials and Methods for details). In Figure 3a, we compared the contours predicted by *VolPy* with manual annotations on three example datasets: *VolPy* is able to identify candidate neurons even in conditions of low signal-to-noise and spatial overlap. In order to quantify *VolPy* performance in detecting neurons, we employed a precision/recall framework (see Material and Methods for details), which accounts for the amount of overlap between predicted and ground truth neurons when assigning matches and mismatches (***Giovannucci et al., 2019***). In Figure 3b and Table 3 we summarize the *F*_1_ score for all the probed datasets. The results indicate that our segmentation approach performs well provided sufficient neurons are fed to train the algorithm. Indeed, *VolPy* obtained *F*_1_ scores of 0.89 ± 0.01 on the L1 dataset (532 neurons in total), 0.71 ± 0.02 on the TEG datasets (107 neurons), and 0.46 ± 0.07 on the HPC dataset (39 neurons). In case of TEG, the performance of *VolPy* is fair considering that the network was trained with only two datasets of this type. In the HPC datasets however the performance on both training and test sets is relatively inferior. We hypothesize that this is due to the fact that not enough data are available (see #neurons column in Table 1), possibly combined with the low signal to noise typical of this dataset type. Note that we used a single neural network trained on the three dataset types simultaneously. Despite clear differences in neuronal shapes, size, SNRs and data acquisition system the network performed well across them, suggesting that it will generalize to similar datasets. However, new datasets deviating substantially from these typologies will need to be added to the training set to improve generalization performance.

**Figure 3.**
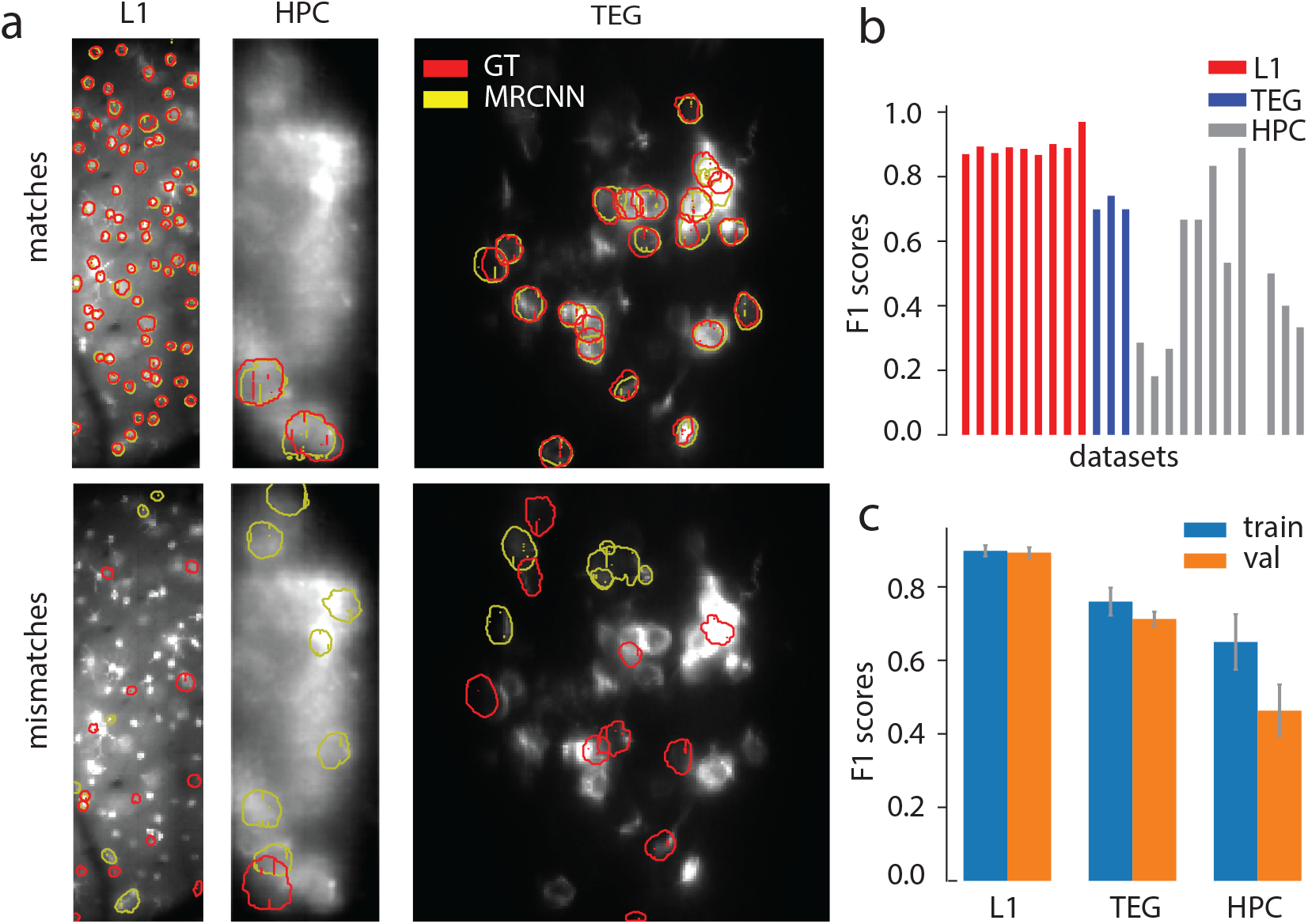
Evaluation of *VolPy* segmentation. (a) Evaluation of segmentation against three manually annotated datasets including mouse sensory cortex (left, Voltron, dataset L1.00.00), mouse hippocampus (center, paQuasAr3, dataset HPC.48.08), and larval zebrafish (right, Voltron, dataset TEG.01.02). In the upper panels, neurons that are found by both *VolPy* (yellow contours) and manual annotators (red contours) are displayed over the mean image. The bottom panels display neurons that are found by *VolPy* but are not present in the ground truth (yellow, False Positives) and neurons that are in the ground truth but are not found by *VolPy* (red, False Negatives). (b) *F*_1_ score performance of *VolPy* for all the evaluated datasets. The *F*_1_ score is computed through stratified cross-validation (see Material and Methods). (c) Average performance on training and validation sets grouped by dataset type (see also Table 3). Error bar represents one standard deviation.

**Table 3.**
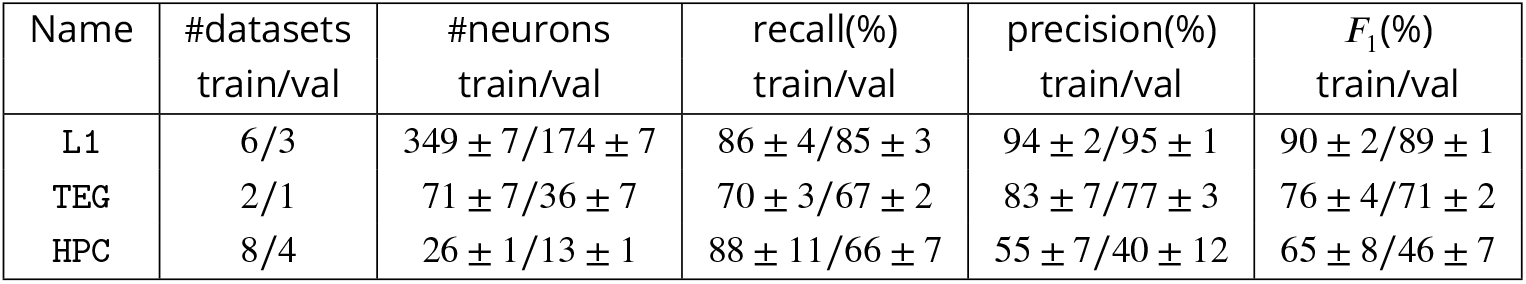
Results of *VolPy* for segmentation. For each type of datasets, number of datasets, number of neurons, recall, precision, *F*_1_ score for training and validation computed by stratified cross-validation are provided.

| Name | #datasets<br>train/val | #neurons<br>train/val | recall(%)<br>train/val | precision(%)<br>train/val | $F_1$(%)<br>train/val |
| --- | --- | --- | --- | --- | --- |
| L1 | 6/3 | $349 \pm 7/174 \pm 7$ | $86 \pm 4/85 \pm 3$ | $94 \pm 2/95 \pm 1$ | $90 \pm 2/89 \pm 1$ |
| TEG | 2/1 | $71 \pm 7/36 \pm 7$ | $70 \pm 3/67 \pm 2$ | $83 \pm 7/77 \pm 3$ | $76 \pm 4/71 \pm 2$ |
| HPC | 8/4 | $26 \pm 1/13 \pm 1$ | $88 \pm 11/66 \pm 7$ | $55 \pm 7/40 \pm 12$ | $65 \pm 8/46 \pm 7$ |

### *VolPy* detects with fidelity single spikes from voltage imaging data

We validated the *VolPy* SpikePursuit algorithm on three voltage imaging datasets in which electrophysiology was simultaneously recorded with voltage imaging (see Figure 4). We automatically analyzed voltage imaging data *in-vivo* recordings from mouse L1 neocortex and Zebrafish Tegmental area (***Abdelfattah et al., 2019***) with the *VolPy* pipeline. The output of the algorithm are spatial footprints, voltage traces, and corresponding spike timings. Spikes for electrophysiology recordings were obtained by thresholding (see Figure 4). Spikes are matched against ground truth by solving a linear sum assignment problem using the Hungarian algorithm (see Material and Methods for details). The *F*_1_ score of each dataset (see Figure 4b) is computed relying on a precision/recall framework based on matched and unmatched spikes. We observe that *VolPy* performs well on all datasets (the *F*_1_ score across three datasets is 0.94 ± 0.03) and confirms that single spikes from voltage imaging data can be automatically extracted with fidelity.

**Figure 4.**
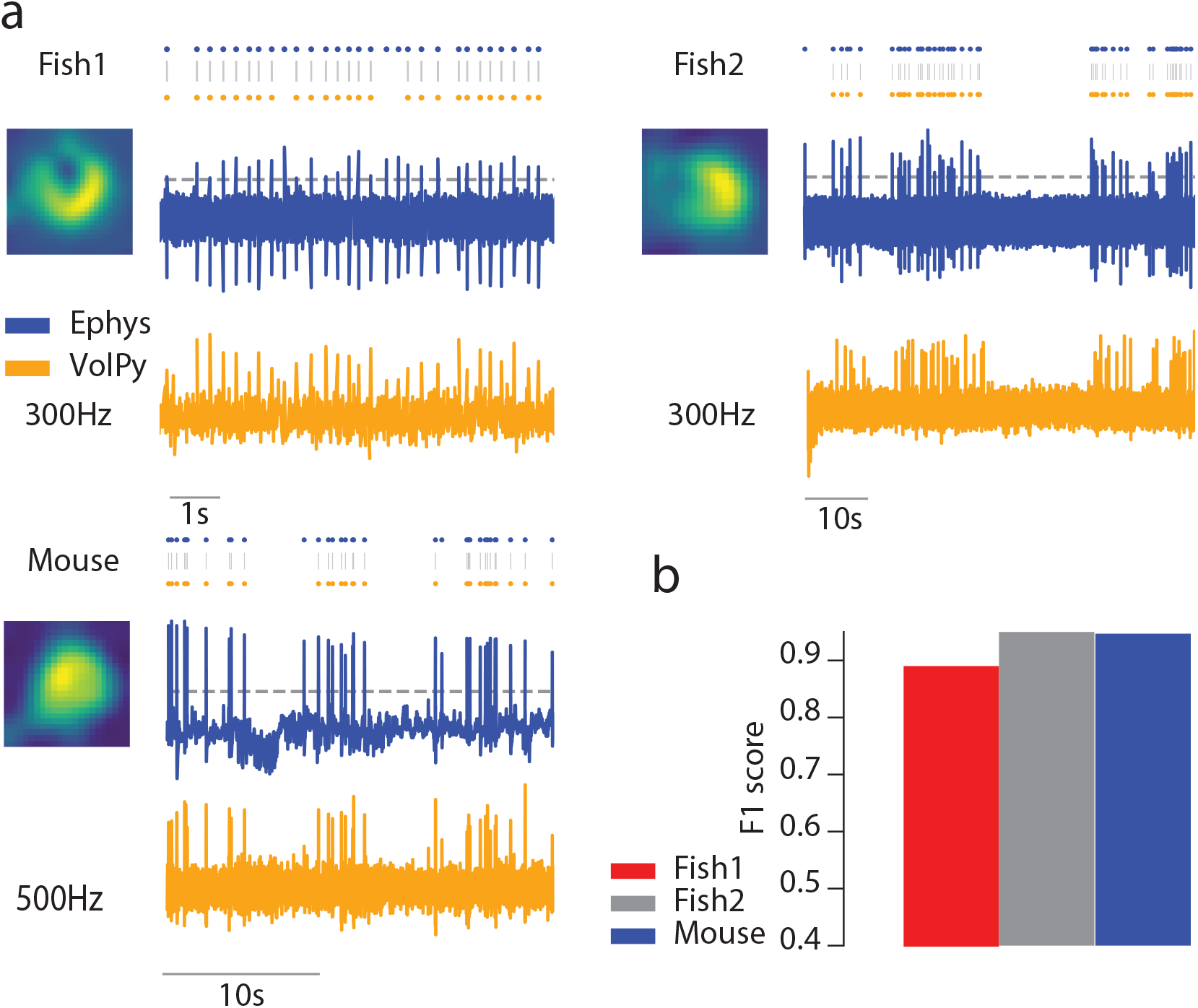
Validation of *VolPy* performance against electrophysiology. (a) Performance of *VolPy* in detecting spikes validated on three datasets from zebrafish (Fish1 and Fish2) and mouse (Mouse). For each dataset, the denoised spatial filter of the target neuron is presented on the left, while electrophysiology (top, blue) and fluorescence signal denoised by *VolPy* (bottom, orange) are reported on the right. Spikes from electrophysiology (blue dots) are obtained by thresholding (gray horizontal dotted line) while spikes from voltage imaging (orange dots) are the output of *VolPy*. Spikes are matched between the two groups by solving a linear assignment problem (see Material and Methods, gray vertical lines). (b) The *F*_1_ scores for each dataset are computed based on the matched and unmatched spikes.

### *VolPy* enables the analysis of large voltage imaging datasets on small and medium sized machines

We examined the performance of *VolPy* in terms of processing time and peak memory for the datasets presented above. We ran our tests on a linux-based desktop (Ubuntu 18.04) with 16 CPUs (Intel(R) Core(TM) i9-9900K CPU @ 3.60GHz) and 64 GB of RAM. For segmentation, we used a GeForce RTX 2080 Ti GPU with 11 GB of RAM memory.

Figure 5a reports the *VolPy* processing time in function of the number of frames. The results show that the processing time scales linearly in the number of frames. Processing 50 candidate neurons in a 1.5 minutes long video (512*128 pixel FOV) takes about 9 minutes. SpikePursuit (red bar) accounts for most of the processing time.

**Figure 5.**
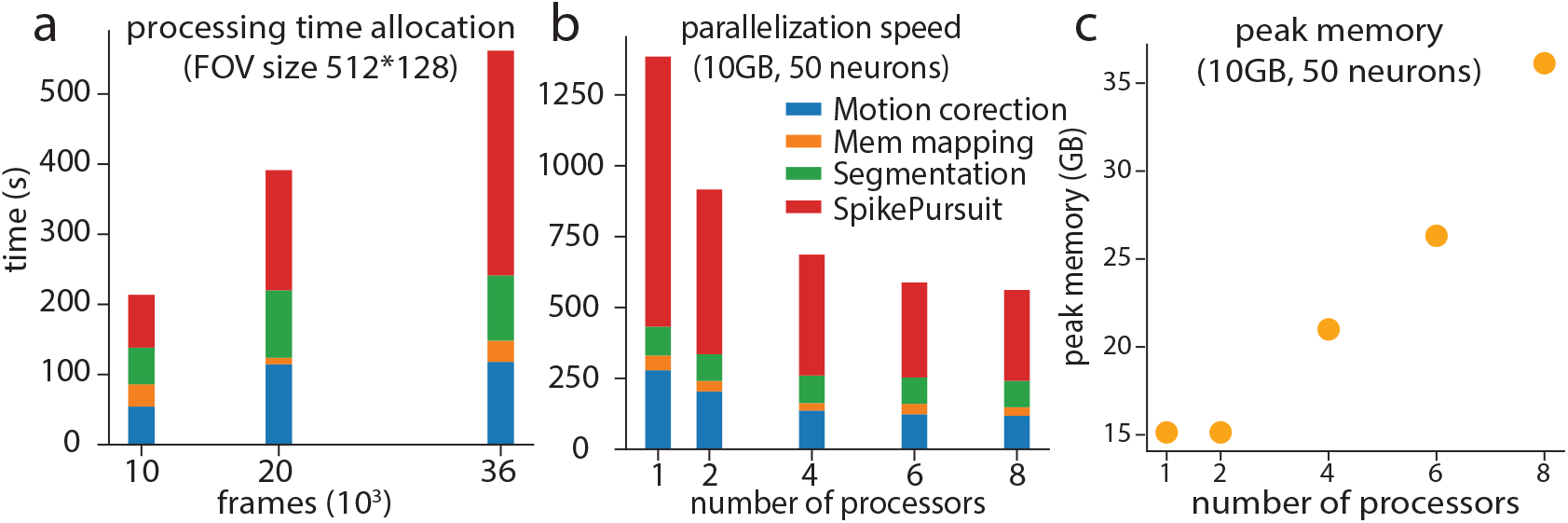
Time and memory performance of *VolPy*. (a) Processing time for *VolPy* as a function of the number of frames on a 512*128 pixels dataset initialized with 50 neurons. Processing time is the sum of motion correction (blue), memory mapping (orange), segmentation (green), and SpikePursuit (red) times. The results indicate a near linear scaling of the processing time with the number of frames. (b) Processing time for *VolPy* utilizing 1, 2, 4, 6 and 8 processors in parallel on a 10 GB dataset and 50 detected neurons. *VolPy* achieves a 2.5-fold speed up when running in parallel on 8 cores. (c) Peak memory usage of *VolPy* in function of the number of processors. Processing in parallel can lead to fair speed gains by regulating the trade-off between time and memory consumption.

In order to probe the benefits of parallelization, we ran *VolPy* 5 times on the same hardware while limiting the runs to 1, 2, 4, 6 and 8 CPUs respectively (Figure 5b). We observed significant performance gains due to parallelization, especially in the motion correction and SpikePursuit phase, with a maximum speed-up of 2.5X. Simultaneously, we recorded the peak memory usage of *VolPy* while running on a different number of CPUs for each run. Figure 5c shows how the peak memory increases with the number of threads. Therefore, *VolPy* enables speed gains by trading-off execution time for memory usage.

## Discussion

### Enabling automated and scalable analysis of voltage imaging data

Recording voltage changes in populations of neurons is necessary to dissect the details of information processing in the brain. Voltage imaging is currently the only technique that promises to achieve this goal with high spatio-temporal resolution. Indeed, voltage imaging has a long history of development, having been widely used for in-vivo studies in the past. However, poor signal-to-noise ratio, photo-toxicity, bleaching, and other difficulties have so far hindered its wider use to answer questions at a cellular level. Recently, however, voltage imaging seems to have reached a point of inflexion and some notable examples are leading the way to new exciting developments (***Abdelfattah et al., 2019***; ***Roome and Kuhn, 2018***; ***Adam et al., 2019***; ***Piatkevich et al., 2019***, ***2018***).

Despite the recent availability of high quality datasets, there is currently no established and validated pipeline for the analysis of voltage imaging data. The unprecedented data size (one order of magnitude larger than already challenging calcium imaging datasets), low SNR, and high degree of signal mixing have so far limited the development of novel algorithms. Moreover, the lack of a universal benchmark prevents further quantitative comparisons. In this paper we provided both a corpus of manually segmented datasets and *VolPy*, the first turn-key, fully automatic, scalable and reproducible pipeline for the analysis of large scale voltage imaging datasets. *VolPy* equips experimenters with efficient computational routines for data handling, motion correction, memory mapping, neuron localization and segmentation, trace denoising and spike extraction. *VolPy* builds upon several optimized and robust routines of the well-established *CaImAn* framework, which it extends to deal with voltage imaging data.

In particular, our contributions develop along the following lines. We provided a corpus of 24 annotated datasets from different brain areas, collection systems and voltage indicators. We developed an automated segmentation supervised algorithm which relies on a Mask R-CNN neural network architecture. We trained a *single* network for all types of considered datasets and evaluated it using cross-validation. The algorithm performance is excellent when enough training data is provided, but smoothly degrades when input data is scarce for specific types of datasets. Regarding trace denoising and spike extraction approaches, we built upon the SpikePursuit algorithm (***Abdelfattah et al., 2019***) and extended it to make it fully automatic, to improve its reproducibility, performance, and to enhance its scalability. We benchmarked the performance of *VolPy* in extracting action potentials against ground truth electrophysiology, with results averaging an outstanding *F*_1_ score of 0.94. Scalability is achieved by leveraging the infrastructure previously deployed in *CaImAn*, which we adapted to enable the parallel processing of multiple neurons. *VolPy* enables a time-memory trade-off which can be tuned based on the available computing power. We demonstrated that *VolPy* enables voltage imaging data analysis on desktop computers. Towards our goal of providing a single package for dealing with standard problems arising in the analysis of imaging data, *VolPy* is fully integrated into *CaImAn* and is therefore immediately available to many laboratories worldwide. The proposed framework is therefore poised to promote the distribution of voltage imaging within the neuroscience community, and in consequence to open the path to a new generation of experiments bridging the gap between electrophysiology and optical imaging.

### Future directions

As more data become available and more users adopt *VolPy*, we plan to develop a graphical user interface for experimentalists to manually label datasets and transfer the resulting annotations to a cloud server, which we will periodically use to retrain and improve the performance of our system.

SpikePursuit is built upon linear methods with a small number of easily-interpreted parameters. An advantage of this approach is that the parameters for can be tailored to different datasets by end users (for example: context area, number of spikes used for templates, filter bandwidth and confidence in segmentation). A continuing challenge for optical physiology is the limited electrophysiological ground truth available for training complex spike detection models. As more training data become available, we expect machine learning approaches to supersede the spatial and/or temporal filtering steps used by SpikePursuit within *VolPy*. Even without large training datasets, algorithmic improvements may be possible. For example, SpikePursuit implements efficient but approximate spike detection using matched filtering with a single template, but could be extended e.g. to include multiple templates or subtractive interference cancellation (***Franke et al., 2015***). *VolPy* and the datasets provided here provide an ideal common ground for comparing such methods.

Finally, and similar to our work in calcium imaging (***Giovannucci et al., 2017***), we plan to generalize our algorithm to real-time scenarios, where activity of neurons needs to be inferred on the fly and frame-by-frame.

## Materials and Methods

### Motion correction & Memory mapping

*VolPy* performs motion correction and memory mapping similarly to *CaImAn* (***Giovannucci et al., 2019***). For motion correction, *VolPy* uses the NoRMCorre algorithm (***Pnevmatikakis and Giovannucci, 2017***) which corrects non-rigid motion artifacts in two steps. First, motion vectors are estimated with sub-pixel resolution for a set of overlapping patches which tile the FOV. Second, the sub-pixels estimates are upsampled to create a smooth motion field for each frame, which is then applied to correct the original frames. Unlike previously, our new implementation enables to perform motion correction on a large number of small files containing a single image (the typical output of fast imaging cameras). This is achieved by multiple parallel processes reading files incrementally and concurrently from the hard drive. This avoids the time- and memory-intensive job of transforming single image files into multi-page or hdf5 files. This modification leads to significant savings in memory, hard drive space and speed.

*VolPy* adopts an optimized framework for efficient parallel data read and write. This framework is based on the ipyparallel and memory mapping Python packages (see ***Giovannucci et al. (2019)*** for more details). In brief, the former enables the creation of distributed clusters across workstations or HPC infrastructures, and the latter enables reading and writing slices of large data tensors without loading the entire file into memory. This is especially important for voltage imaging, considering the larger file sizes compared to calcium imaging. This framework, which in *VolPy* implements an embarrassingly parallel infrastructure, is used across different steps of the pipeline:
 
- The output of the motion correction operation is saved into a set of F ordered Python memory map files without creating any other intermediate files. This is done in parallel over all the processed movie chunks.
- The motion corrected F order files are then consolidated into a single C ordered memory map file. This is also performed in parallel over many processes.
- During trace denoising and spike extraction, each process loads and processes in parallel a small portion of the field of view.

### Creating a corpus of annotated datasets

We generate a corpus of annotated datasets in which neurons are manually segmented. For neurons to be selected at least one of the following criteria needed to be met: (i) Both annotators had to agree on the selection; (ii) Neurons were very clear on at least one of the three cues; (iii) Neurons were moderately clear in one of the summary images and appeared clearly in a few frames of the local correlations movie.

Figure 8 shows the process of selecting neurons. Ground truth is inferred by human labelers from mean and local correlation images as well as a local correlation movie which highlights active neurons. Relying on these three visual cues, two annotators marked the contours of neurons (yellow color) using the ImageJ Cell Magic Wand tool plugin (***Walker, 2014***) and saved the result into the ROI manager in ImageJ.

### Segmentation via convolutional networks

*VolPy* uses a variation of the Mask R-CNN framework (see Figure 6) to initialize spatial footprints of neurons. In the following section we will introduce the Mask R-CNN framework in the *VolPy* context.

**Figure 6.**
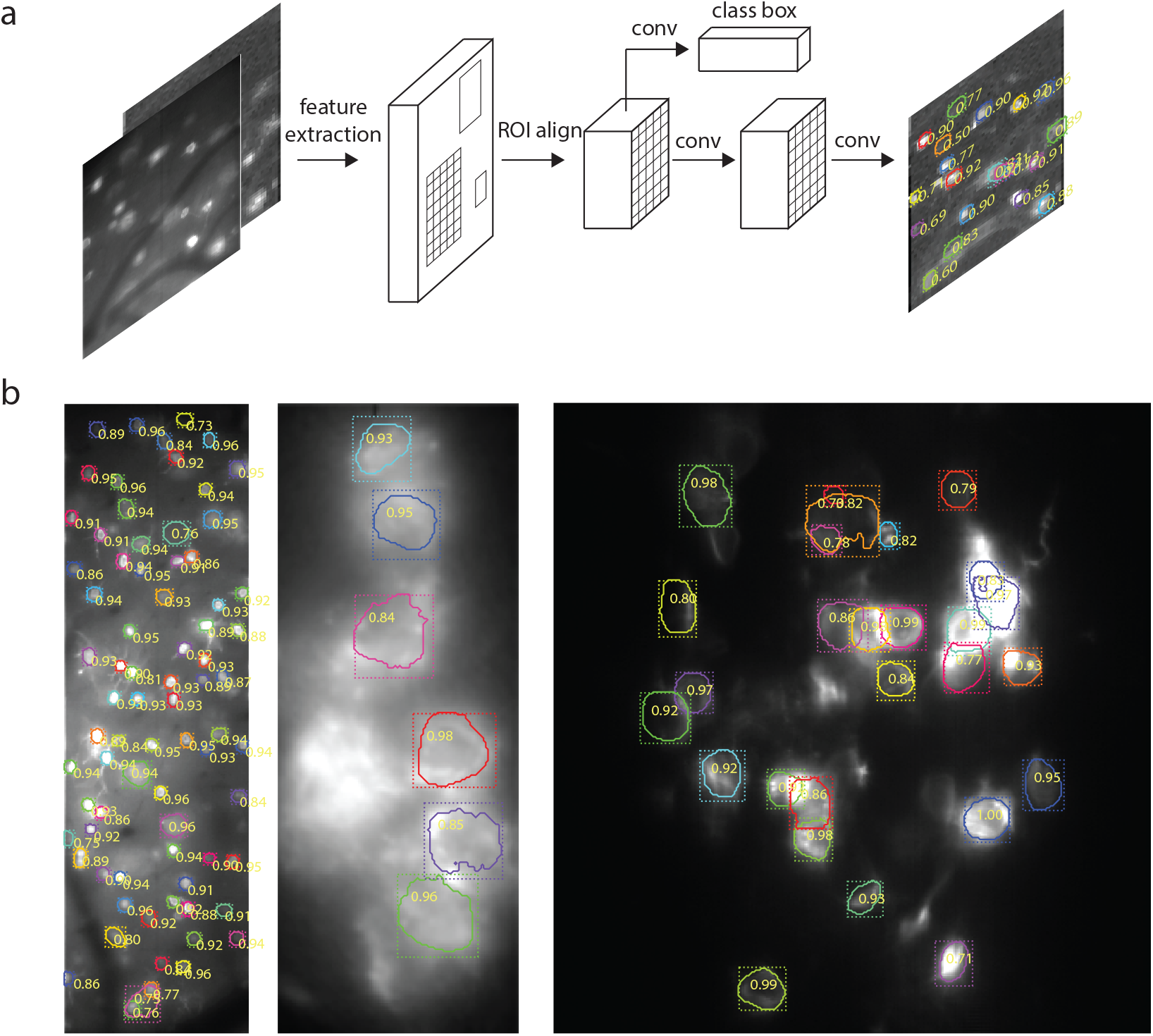
Segmentation algorithm of *VolPy*. (a) Mask R-CNN framework for segmenting neurons with summary images as input to the network. (b) The output of *VolPy* segmentation algorithm run on three example datasets from sensory cortex (left, Voltron, dataset L1.00.00), mouse hippocampus (center, paQuasAr3, dataset HPC.48.08), and larval zebrafish (right, Voltron, dataset TEG.01.02). The mean images are overlaid with contours (solid line), bounding boxes (dotted line) and detection confidence for each candidate neuron. Only neurons with detection confidence greater than 0.7 are displayed.

#### Mask R-CNN

Mask R-CNN (Figure 6a) is a network architecture which provides simultaneous object localization and instance segmentation via a combination of two network portions: backbone and head. The backbone features a pre-trained convolutional network (such as VGG, ResNet, Inception or others) for extracting features of the input image. Mask R-CNN also exploits another effective backbone: Feature Pyramid Networks (FPN) (***Lin et al., 2017***), a top-down architecture with lateral connections, which enables the network to extract features on multiple scales from the feature maps. In the head, based on the extracted features, a Region Proposal Network proposes initial bounding boxes for each candidate object, which are fed to two downstream branches. One of them is trained to predict a class label and a bounding box offset which refines the initial bounding box, while the other branch outputs a binary mask for each candidate object.

#### *VolPy* Mask R-CNN

We adapt Mask R-CNN to our purpose by introducing the following modifications. We choose a combination of ResNet-50 pre-trained on the COCO dataset and FPN as the backbone. The input of the network is a three channel image: two for the mean images and one for the correlation image. The three channel image is necessary in order to re-use the first few layers which were pre-trained on the COCO dataset. The network is trained to predict only one class, neuron or not neuron (background) instead of a multi-label output.

##### Training

We randomly crop the input image into 128×128 crops and apply the following data augmentation techniques using the *imgaug* (***Jung et al., 2019***) package: flip, rotation, multiply (adjust brightness), Gaussian noise, shear, scale and translation. Each mini-batch contains six cropped images. We train on one GPU the heads (the whole network except the ResNet) of the network for 2k iterations with learning rate 0.01 and then train layers after the first three stages of the ResNet (28 layers) for another 2k iterations with learning rate 0.001. We use stochastic gradient descent as our optimizer with a constant learning momentum 0.9. The weight decay is 0.0001.

##### Validation

Images are padded with zeroes to make width and height multiples of 64 so that feature maps can be smoothly scaled for the Feature Pyramid Network. We only choose neurons with confidence level greater or equal to 0.7.

### Trace denoising and spike extraction

Trace denoising and spike inference are performed by an improved version of the SpikePursuit algorithm (***Abdelfattah et al., 2019***), in which we optimized for speed, memory usage, and accuracy. The pseudo-code for the associated computational steps is reported in Algorithm (1) and Figure 7. The algorithm starts by approximating a neuronal signal and the background contamination from the ROI provided by the segmentation step. The algorithm then proceeds iteratively to detect the most prominent spikes, extract a waveform template from detected spikes, use the template to recover similarly-shaped spikes, reconstruct the trace from the recovered spikes, and use the reconstructed trace to improve the spatial filter. These steps are explained in more details below.

**Figure 7.**
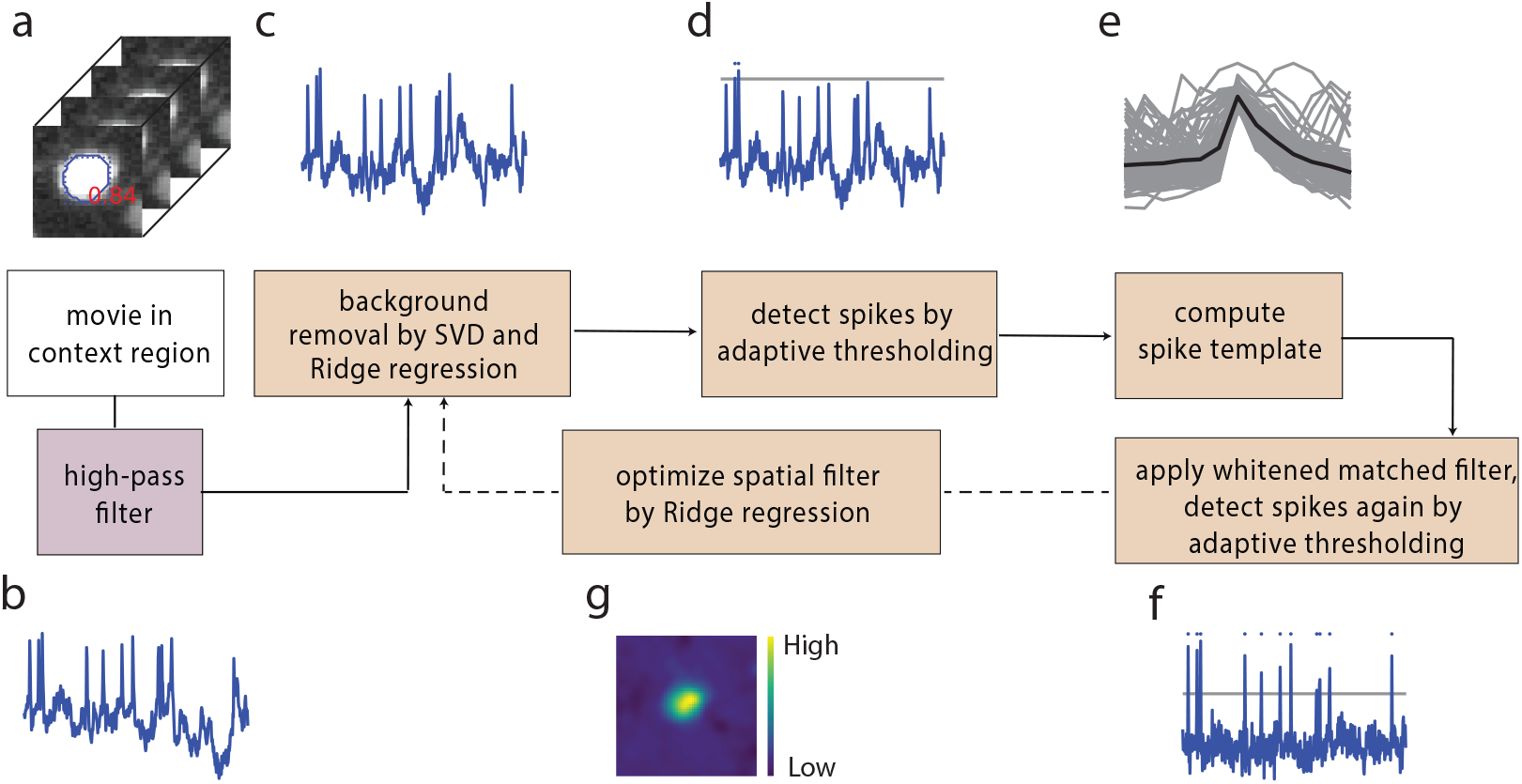
Algorithm for fluorescence trace denoising and spike inference. (a) A small section of the movie (context region) encompassing a candidate neuron and a neighboring area is loaded from the C ordered memory mapped file. (b) After high-pass filtering the movie, the initial temporal trace of the candidate neuron is approximated either from the mean signal of the ROI region pixels, or by applying the spatial filter to the context region if an initial spatial filter is provided. Afterwards, two big steps are executed in loop until convergence (or a maximum of 5 steps). The former ((c),(d),(e) and (f)) estimates spike times, and the latter ((g)) refines the spatial filter. (c) We extract the first 8 principal components of the background pixels using singular value decomposition and then remove the background contamination via Ridge regression. (d) After high-pass filtering the trace, we select spikes with peak larger than an adaptively selected threshold (gray dotted line). The total number of peaks detected in the first round is constrained between 30 and 100. Later rounds of spike detection include all spikes. (e) Waveforms of these spikes (gray) are averaged to obtain a spike template (black line). (f) A whitened matched filter is used to enhance spikes which have a similar shape to the template. (g) Refine spatial filter through Ridge regression. Calculate the weighted average of movie (using the refined spatial filter) as the new temporal trace for the next iteration.

#### ROI loading and preprocessing

As a result of segmentation, each candidate neuron has an associated binary mask which represents its spatial extent (ROI region R). The ROI is dilated to get a larger region (50×50 pixels by default) centered on the neuron (context region C). Background pixels are defined as all the pixels in the context region at least *n*_*B*_ pixels (12 pixels by default) away from the ROI region (background region). As a first step, all pixels in the context region are efficiently retrieved from the memory mapped file and high-pass filtered as *Y*_*h*_ to compensate for photo-bleaching (Figure 7a-b, Algorithm 1 lines 1–8). The initial temporal trace **t**_**0**_ associated to a neuron can be approximated either from the mean signal of the ROI region pixels, or as a weighted average across all pixels in the context region when a spatial filter **w** calculated from previous chunk of data was available (Algorithm 1 lines 9–13):

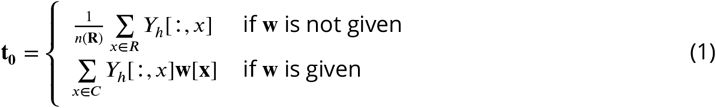

where *n*(*R*) represents number of pixels in the ROI region.

Afterwards, two steps are executed in loop until convergence (or a maximum of 5 steps). The former tries to estimate spike time, and the latter tries to approximate the spatial filter.

#### Spike time estimation

In order to estimate spike times from the fluorescence traces (Figure 2c-f) we proceed as follows. First, we compute the singular value decomposition of the background pixels *Y_b_*:

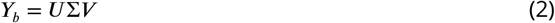

where *U* contains the temporal components. This is then used to remove background contamination via Ridge regression, in which *U_b_*, the first 8 components of *U* is the regressor and the temporal trace is the predictor (Algorithm 1 line 15–16).

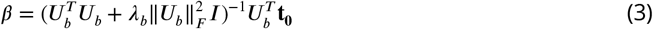

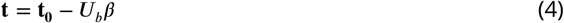

We experienced that a very high signal-to-noise ratio neuron with large spatial footprint included in the background pixels led to poor performance due to unregularized linear regression used at this stage in the original SpikePursuit implementation. Use of non-regularized regression to remove the background can allow real signal to be subtracted from neuron traces if the neuron’s trace is captured by the background PCs. To ameliorate this issue, we modified the original algorithm by adding an *L*_2_ regularizer to penalize large regression coefficients. This provided more reliable results with respect to the original implementation on multiple datasets.

After background removal, the trace is high-pass filtered with a cut-off frequency of 60 Hz and two rounds of spike detection are performed. The first round selects spikes with peak larger than an adaptively selected threshold, while keeping the total number of peaks between 30 and 100 (Algorithm 3). A spike template **z** is computed by averaging all the peak waveforms:

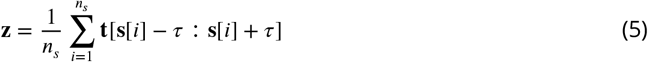

where **s** is the list of spike time, *n*_*s*_ is total number of spikes, *τ* is the half size of window length. Subsequently, a whitened matched filter (***Franke et al., 2015***) is used to enhance spikes with shape similar to a template. More in details, we use the Welch method to approximate the spectral density of the noise in the fluorescence signal. Second, we scale the signal in the frequency domain to whiten the noise. Finally, we convolve with a time-flipped template. The template we used is the peak-triggered average.

The latter round of spike detection incorporates all the spikes detected by applying a newly computed threshold. Then, a reconstructed and denoised trace is computed by convolving the inferred spike train (**q**) with the waveform template:

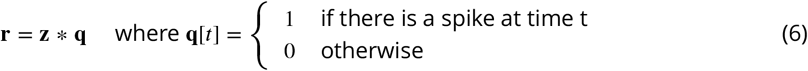

#### Spatial filter refinement

The second step, illustrated in Figure 2g, is to refine the spatial filter. The updated spatial filter is computed by Ridge regression, where the reconstructed and denoised trace is used to approximate the high-passed video (Algorithm 1 line 18):

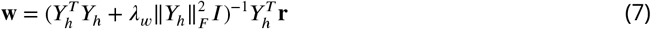

Subsequently, the weighted average of movie with the refined spatial filter is used as the updated temporal trace for the following iteration:

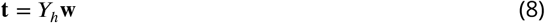

The ridge regression problem was originally solved in SpikePursuit by directly calculating the analytical solution (normal equation). However, the multiplication and inverse of large matrices was computationally inefficient. We decided to apply an iterative and much more efficient algorithm to solve the regression problem (***Paige and Saunders, 1982***) implemented in the Scikit-Learn package (‘lsqr’).

### Precision/Recall Framework to measure segmentation performance

In order to measure the performance of *VolPy* segmentation, we compared the spatial footprints extracted by *VolPy* with our manual annotations (see (***Giovannucci et al., 2019***) component registration for a detailed explanation). In summary, we computed the Jaccard distance (the inverse of intersection over union) to quantify similarity among ROIs, and then solved a linear assignment problem with the Hungarian algorithm to determine matches and mismatches. Once these were identified, we adopted a precision/recall framework and we defined True Positive (TP), False Positive (FP), False Negative (FN), and True Negative (TN) as follows:

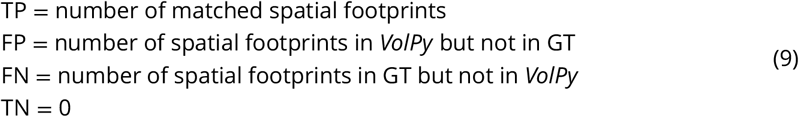

Next we computed precision, recall and *F*_1_ score of the performance in matching as the following:

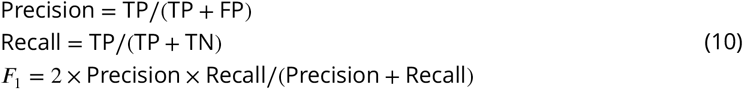

Note that the *F*_1_ score is a number between zero and one. The better the performance of matching, the higher the F1 score.

### Cross-Validation to evaluate segmentation model on limited datasets

In order to decrease the selection bias originated from the separation in training and validation datasets and better evaluate Mask R-CNN model on our limited datasets (24 in total), we performed a stratified three-fold cross-validation. The reason we used a stratified three-fold cross-validation rather than a normal three-fold cross-validation is that we want our model train and validate on each type of datasets. We partitioned datasets into three groups so that arbitrary type of data (L1, TEG, HPC) is partitioned equally into three groups without repetition (Figure 2 train/val column shows one group of the partition). During cross-validation two groups were used as training sets while the remaining one as validation set. The cross-validation process was repeated three times with each group used exactly once as validation set.

For each run of the cross-validation process, we trained a single network and tested it on both training and validation sets. We then computed the mean and standard deviation of the F1 score for different types of datasets with training and validation sets treated separately.

### Spike matching

In order to validate fidelity of spike extraction algorithm, we needed to match spikes extracted from voltage imaging and electrophysiology datasets. Let *v*_1_ *, v*_2_ *, …, v*_*n*_ be the spike time extracted from voltage imaging traces, and *s*_1_ *, s*_2_ *, …, s*_*m*_ be the spike times from electrophysiology ground truth, where n and m are the total number of spikes respectively. We formulate the problem as a linear sum assignment problem. Let **D** be a distance matrix where *D*[*i, j*] is the cost of matching spikes *v*_*i*_ and *s*_*j*_. When the difference of spike-times is larger than a threshold *t*, we assign a large distance value *M*:

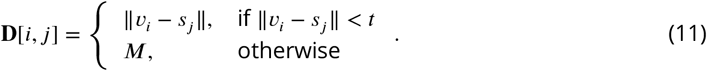

Let **X** be the Boolean matrix where **X**[*i, j*] = 1 if *v*_*i*_ and *s*_*j*_ are matched and 0 otherwise. Each spike can be matched at most once, i.e. at most one element for each row (or column) of **X** can be one. The optimal assignment has the cost:

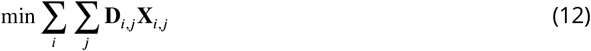

We solve this optimization problem using the Hungarian algorithm implemented in the Scipy package and delete matched spikes whose costs are equal to M. After identifying matches and mismatches, we proceeded similarly to what explained above to extract the *F*_1_ score. We define TP, FP, FN, TN similar to Equation 9:

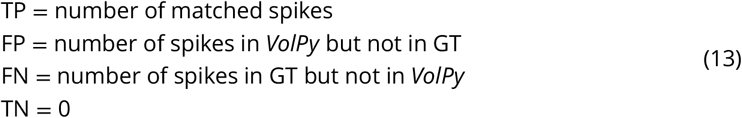

Then we calculated *F*_1_ score same as Equation 10.

## Supporting information

Movie 1. Raw and Correlation image

## Acknowledgments

We thank K Svoboda, A Singh, M Ahrens, T Kawashima, Y Shuai, A Cohen, M Xie for providing voltage imaging datasets. We thank H Eybposh for useful discussions. We thank A Singh for the initial version of Python code, and for insightful comments and discussions.

## Description of Supplemental Movies

**Video 1.** Example of voltage imaging data on mouse neocortex data. Left: Raw data. Right: Local correlation video.

## Description of Supplemental Images

### Algorithmic Details

In the following section we present the pseudocode for several of the routines introduced and used by *VolPy*. Note that the pseudocode descriptions do not aim to present a complete picture and may refer to other work for some of the steps.

**Algorithm 1.**
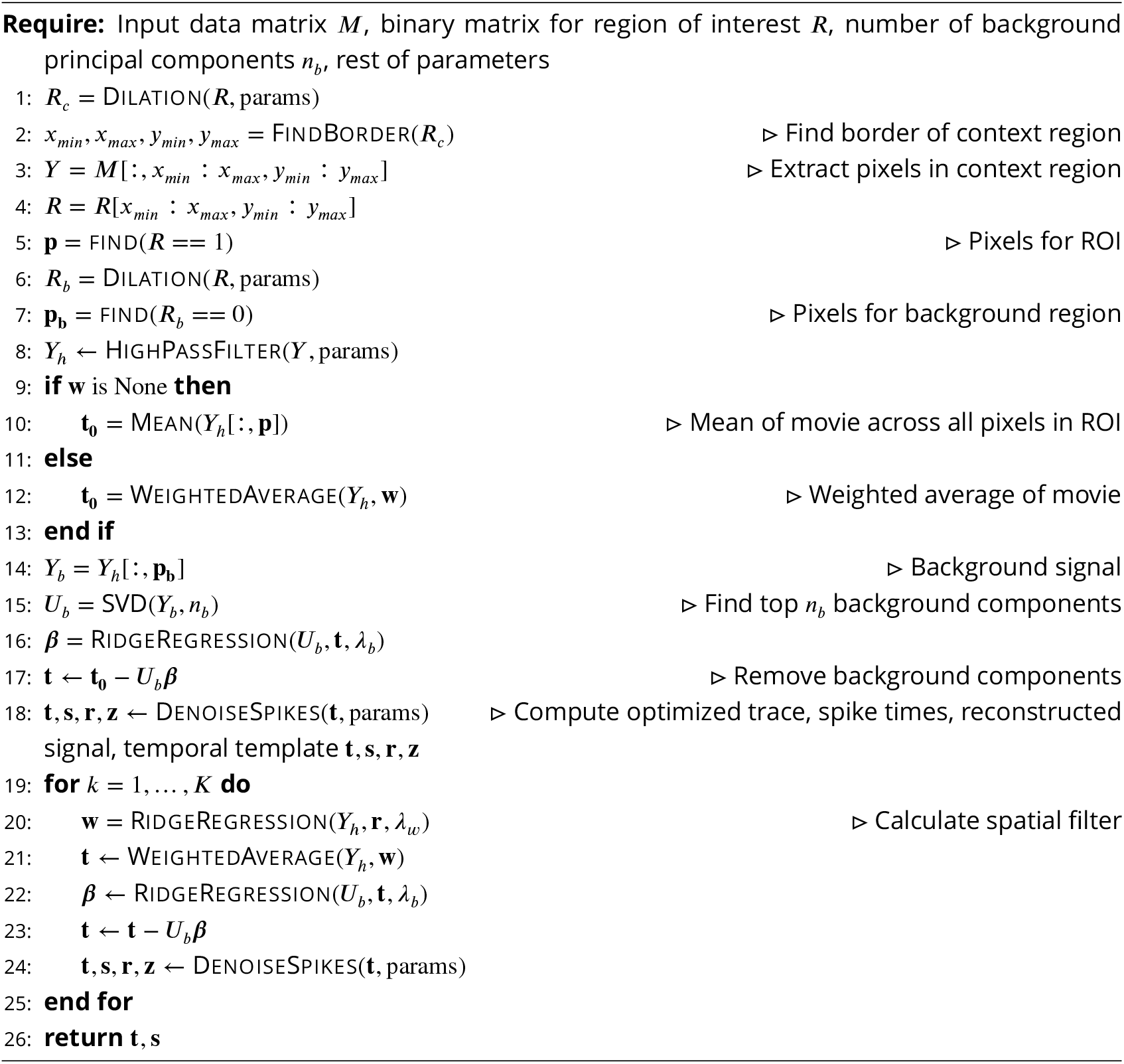
SPIKEPURSUIT

**Figure 8.**
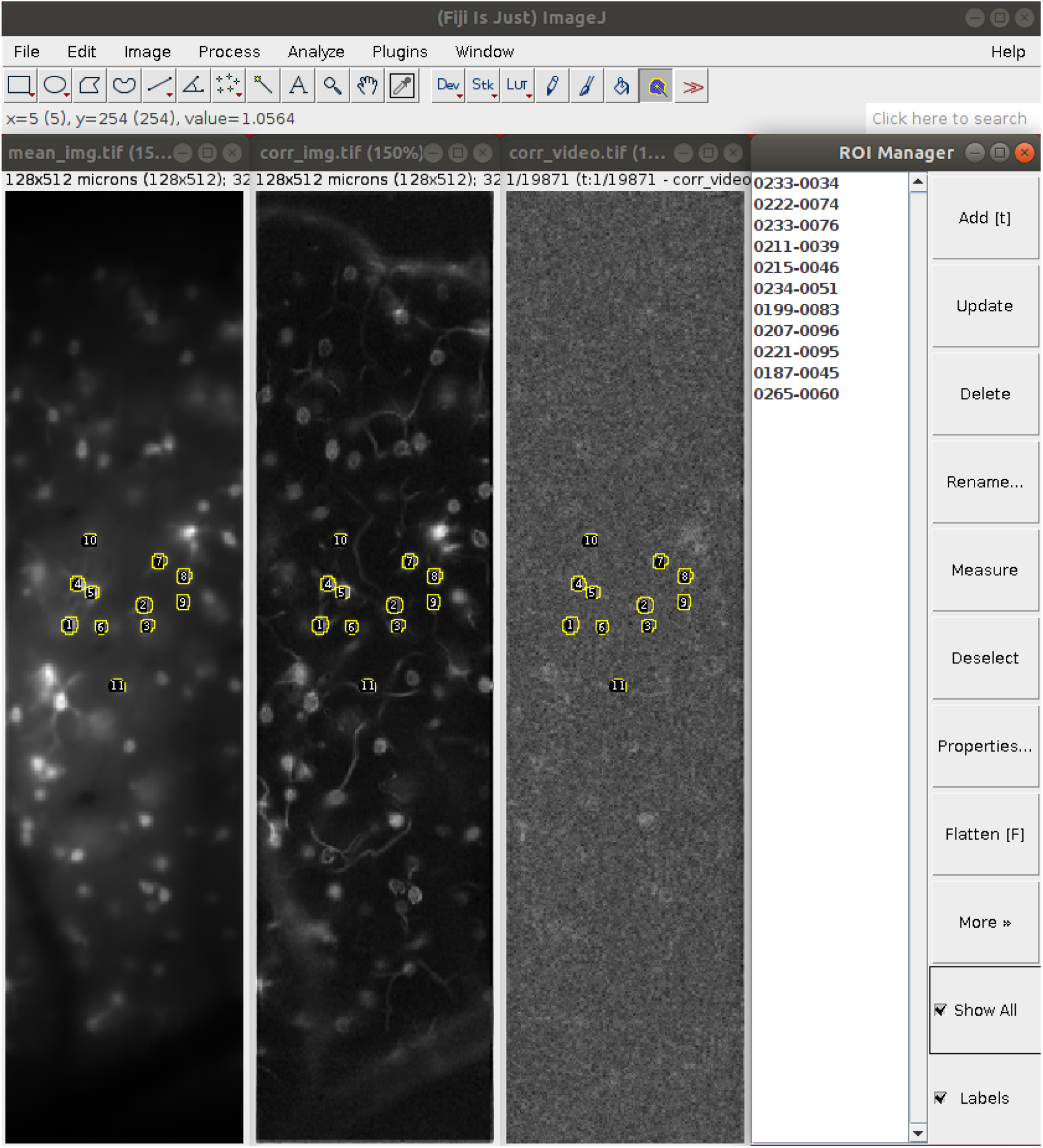
Create manual annotations of voltage imaging datasets with ImageJ. We selected neurons based on mean image (left), correlation image (mid-left) and local correlation movie (mid-right). Two annotators marked the contours of neurons using ImageJ Cell Magic Wand tool plugin and saved selections in ROI manager (right).

**Algorithm 2.**
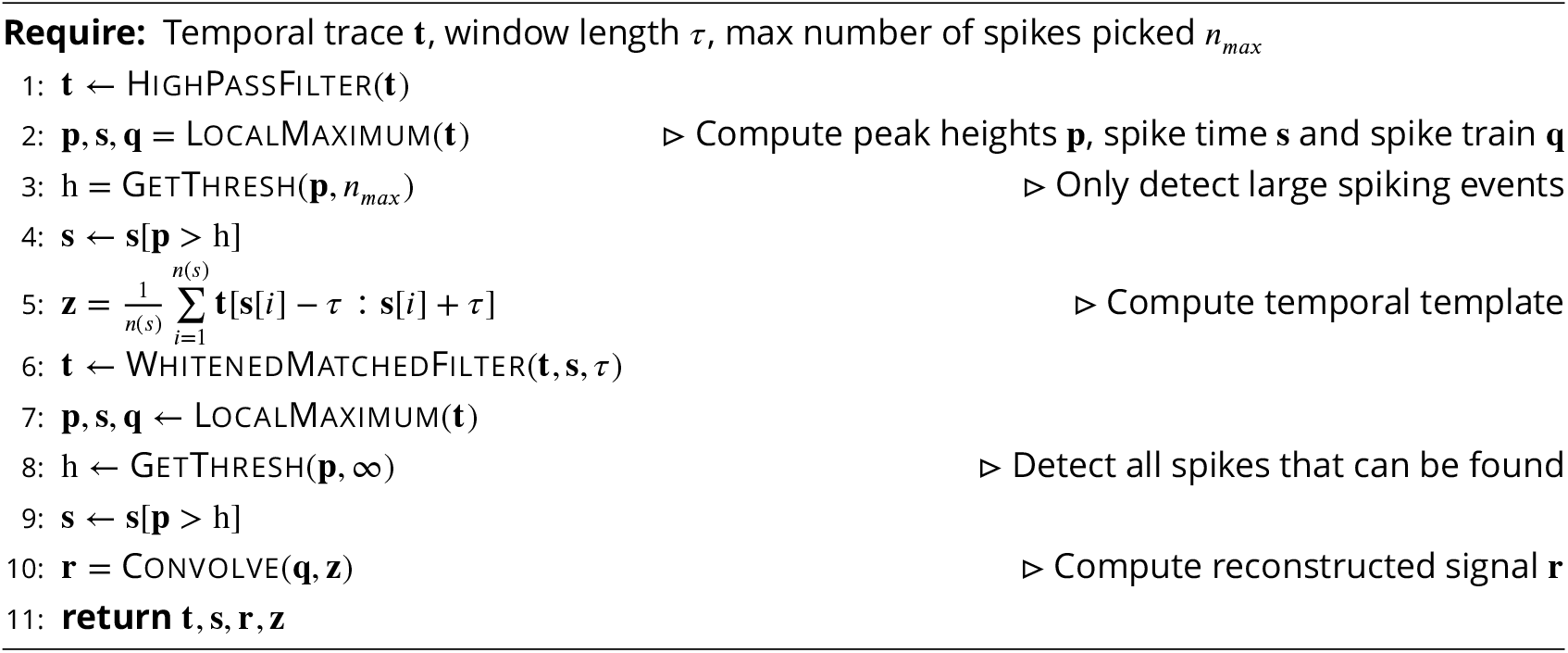
DENOISESPIKES

**Algorithm 3.**
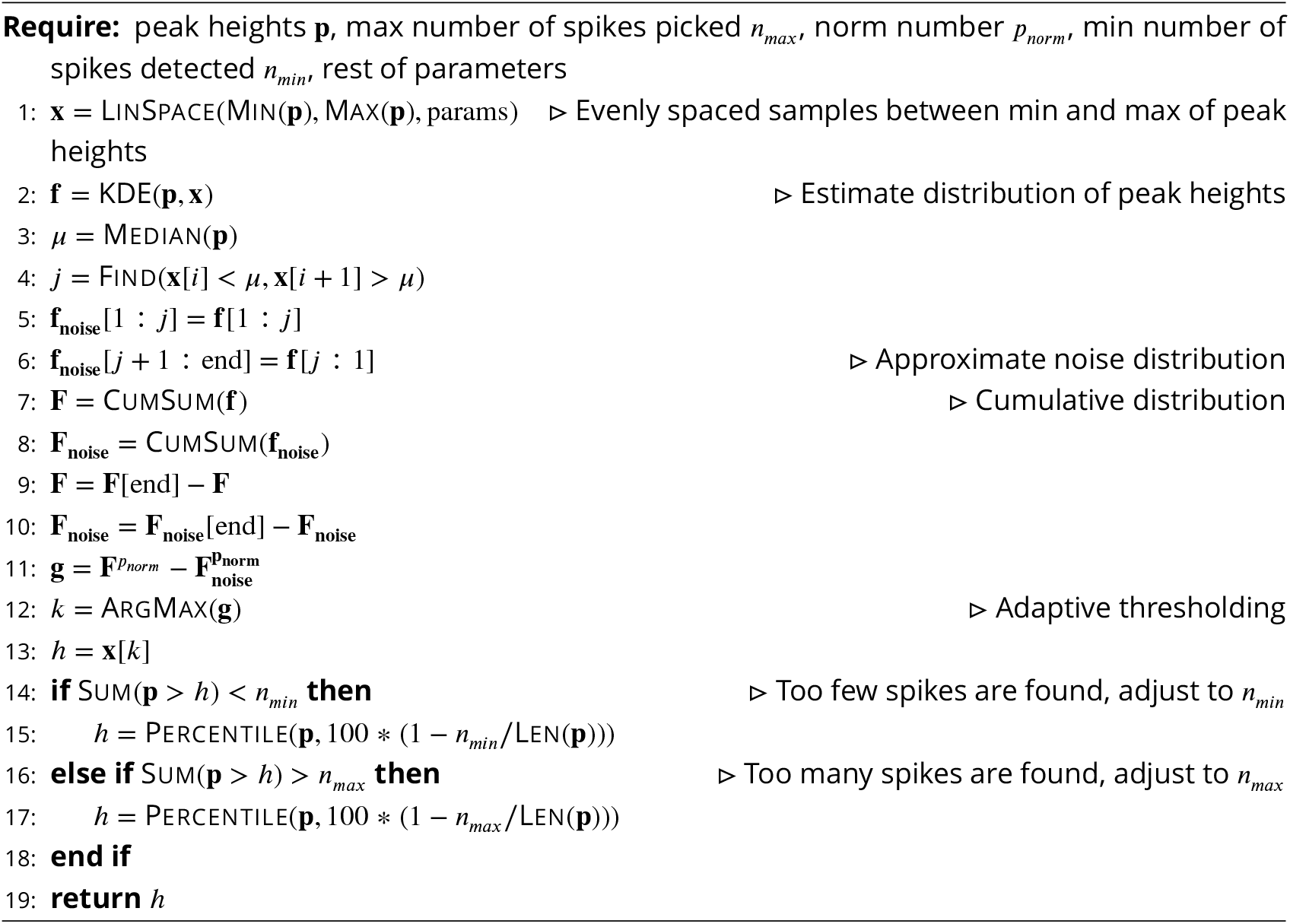
GETTHRESH

**Algorithm 4.**
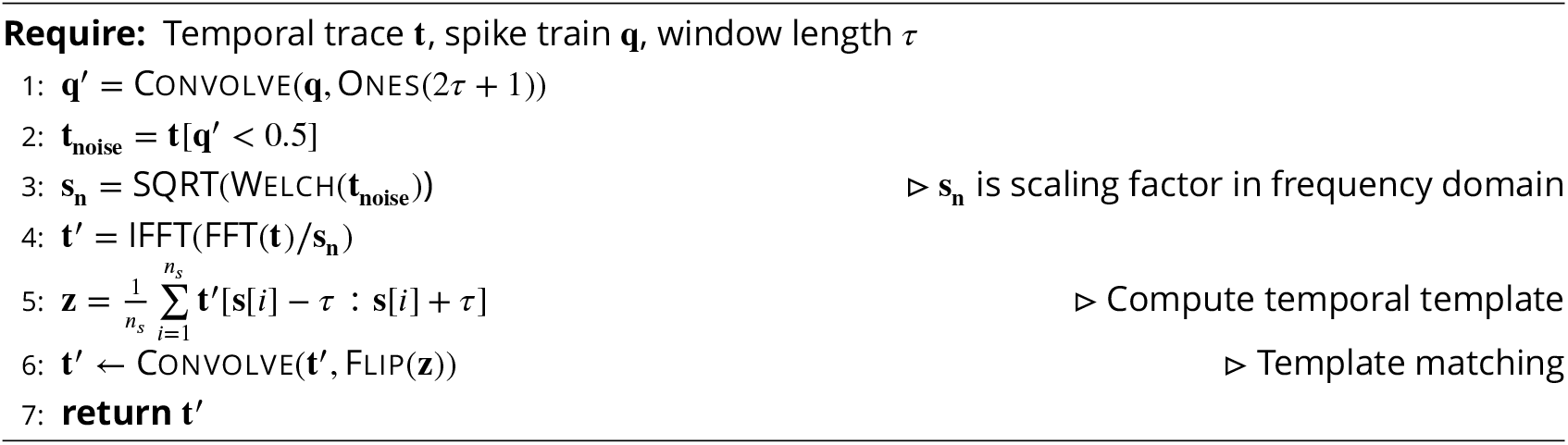
WHITENEDMATCHEDFILTER

